# Assessing analysis methods of brain synchrony in social interaction: Simulation-based comparison

**DOI:** 10.1101/2025.05.12.653588

**Authors:** Satoshi Morimoto, Yasuyo Minagawa

## Abstract

Inter-brain synchrony is an essential measure for investigating social interactive behaviour via hyperscanning. While functional near-infrared spectroscopy is a unique modality for measuring this index in dynamic, real-world interactions, methods to adequately assess inter-brain relationships have not been firmly established, and the overall picture remains unclear. Consequently, in this article, we first briefly summarize analysis methods for examining social interaction by dividing them into static and dynamic measures. Among these, we focus on measures of synchrony and their assessment in correlating behaviours, conducting a simulation-based comparison and analysis. Specifically, we directly compared static and dynamic variants of wavelet transform coherence (WTC), Pearson’s correlation coefficient (CC), and phase mutual information (pMI) using a real fNIRS dataset. Results showed a significant divergence between WTC and CC, while WTC and pMI exhibited similar patterns as static measures. Overall, WTC was suggested to better identify synchrony due to its non-linear, instantaneous, and robust nature. For the latter part, based on other simulation analyses, we propose a new method using generalized linear model (GLM) regression. Simulations with synthetic fNIRS data supported the effectiveness of our proposed method, which can capture the dynamic relationships between inter-brain synchrony and behaviour, even during free interaction.

## Introduction

Functional near-infrared spectroscopy (fNIRS) is a non-invasive brain recording system that can measure changes in oxygenated and deoxygenated haemoglobin. Its low cost, silence, portability, and fewer physical constraints allow us to experiment in a more realistic environment, including naturalistic interaction (Quaresima and Ferrari, 2016). Hyperscanning, the simultaneous recording of brain activity in two or more participants, is a valuable method for directly assessing the relationship in neural activity between participants (Montague *et al*., 2002; Scholkmann *et al*., 2013; Babiloni and Astolfi, 2014; Minagawa *et al*., 2018). Due to its greater resistance to motion artefacts compared to other neuroimaging modalities like electroencephalography (EEG), fNIRS has been successfully used to reveal significant neural synchronisation between dyads during interaction (Czeszumski *et al*., 2020). Ensure that neural synchronies obtained in different measurement systems reflect different physiological aspects of brain activity; similar to functional magnetic resonance imaging (fMRI), fNIRS measures a haemodynamic change that is not a direct measure of neural activation, in contrast to direct electrophysiological measures obtained by EEG and magnetoencephalography (MEG). Therefore, the optimal analysis method should be explored to better fit the nature of the signals and experimental purposes depending on the measurement system. While EEG hyperscanning has accumulated more knowledge and practice, including data analysis methods with its longer history, fNIRS hyperscanning cannot directly employ EEG analysis methods due to differences in temporal resolution. This is a critical point in analysing the relationship between brain signals (e.g., oxygenated haemoglobin) and behaviour (e.g., eye gaze) in the context of social interaction. However, it is possible that fNIRS hyperscanning can still utilise some analysis methods employed in EEG studies, such as mutual information. Taken together, analysis methods for fNIRS hyperscanning are still an emerging field to be explored and clarified.

For fNIRS hyperscanning, the major methods for quantifying inter-brain synchrony are wavelet transform coherence (WTC; Torrence and Compo, 1998; Grinsted *et al*., 2004) and correlation coefficient (Hakim *et al*., 2023). WTC quantifies coherence in the time and scale (i.e., frequency) domain, making it suitable for representing temporal synchrony within specific frequency bands. On the other hand, Pearson’s correlation coefficient (CC) evaluates the linear relationship between two signals. In most studies, WTC was averaged over the time series and frequency ranges of interest (tsWTC). Both the resulting scalar values, tsWTC and CC with band-pass filtered signals, were used as measures of inter-brain synchrony. However, the aspect of inter-brain synchrony being quantified probably differs for each measure (e.g., CC assuming linearity, WTC not assuming linearity for fNIRS data). To the best of our knowledge, no studies have addressed the differences among measures and rationally selected a measure from them based on the research purpose. It is important to explore the characteristics of each measure and its parameter settings for fNIRS signals, and to present their differences as measures of inter-brain synchrony.

Another problem to be raised here is synchrony analysis in relation to behavioural correlations. For example, during free interaction, inter-brain synchrony is likely to vary along the time course depending on the magnitude of the interaction. Combined analysis using both behavioural information and brain activity is necessary in this case. However, behavioural association analysis for dynamic inter-brain synchrony is not well established, despite it being one of the most important challenges for social interaction studies.

Although social interaction studies with fNIRS have employed different analysis methods to assess inter-brain synchrony, the relationships between these methods and their advantages and disadvantages remain unclear, as stated above. Accordingly, in this paper, after briefly summarising the overview of analysis methods for hyperscanning data, we conducted simulations to examine the quantitative comparison of different inter-brain synchrony measures of fNIRS signals. We illustrated the characteristics of each measure and highlighted the differences between them. Two sets of simulations were performed: one for comparisons between static measures and the other for comparisons between dynamic (i.e., temporally varying) measures. Furthermore, in the latter part, we focus on how to correlate behavioural data with the fNIRS synchrony signal, which is relatively slow. Specifically, by evaluating the behavioural relevance of the dynamic measures, we propose a new method using the WTC-GLM approach and demonstrate its performance through simulations. Finally, we discuss the future directions for the quantification of fNIRS hyperscanning data.

### Analysis Methods for fNIRS Hyperscanning data

This section provides an overview of the analysis methods for fNIRS data measured in the hyperscanning setting (Table 1). We categorise measures for quantifying neural synchrony into two distinct groups: one consisting of metrics averaged over specific experimental conditions (i.e., static measures), and the other consisting of metrics that are time-varying (i.e., dynamic measures). In accordance with this categorisation, we make direct comparisons of the methods within each group in the following section.

**Table. 1.**
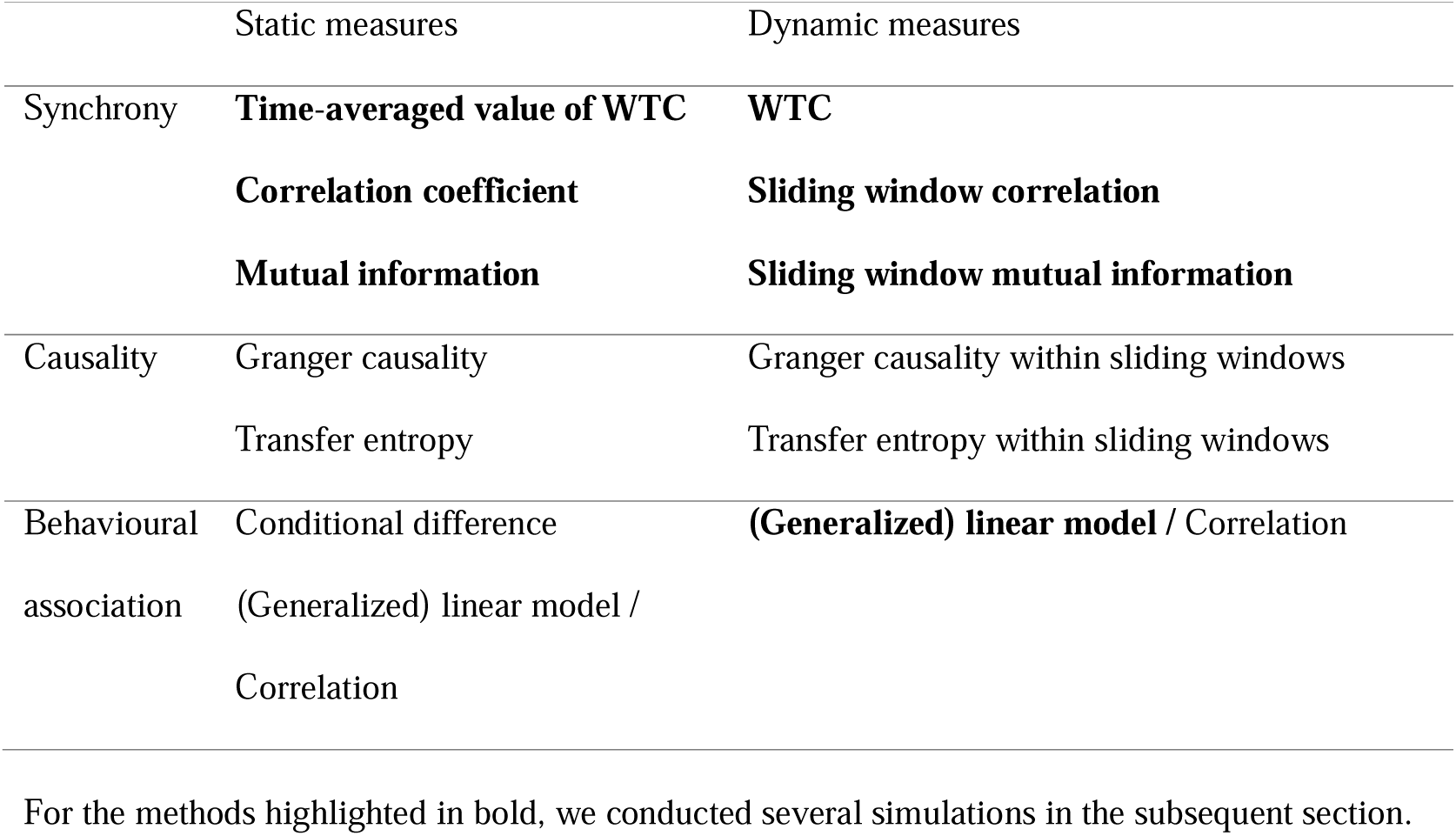
Example of methods for analysing the hyperscanning fNIRS data.

Causality is another metric used to quantify inter-brain relationships (Hakim *et al*., 2023), including Granger causality (Seth *et al*., 2015) and transfer entropy (Vicente *et al*., 2011). Although we do not provide detailed information on the causality analyses for the purpose of this article, we discuss the future direction of causality analysis in the Discussion section. Similar to the synchrony measures, we categorise the causality measures in the same manner.

Methods for behavioural associations are also categorised into two groups. In this study, we focused solely on associations with inter-brain synchrony.

We briefly introduce each measure of inter-brain synchrony and behavioural association with the measures in turn. In addition, we provide a review of the fundamental techniques for preprocessing the fNIRS signal, which is directly related to the synchrony measures.

#### Preprocessing

Preprocessing is the most important part of fNIRS signal analysis. After obtaining the raw light intensity data, the optical density is first calculated using a logarithmic transformation, then correct the motion artefact, apply filtering to remove non-cortical physiological signals, and finally convert it into the concentration changes of oxygenated haemoglobin (Δoxy-Hb) and deoxygenated haemoglobin (Δdeoxy-Hb) using the modified Beer-Lambert law (see Yücel *et al*., 2021 for details). fNIRS is relatively robust to motion artefacts compared to EEG, however, changes in optode placement can cause significant artefacts. For data collected during realistic interactions, correction of motion artefacts is crucial for obtaining reliable results (Cooper *et al*., 2012; Pinti *et al*, 2019; Gemignani and Gervain, 2021; Huang *et al*., 2022).

The primary physiological noise components in fNIRS include the heartbeat (∼1 Hz), respiration (∼0.3 Hz), Mayer waves (∼0.1 Hz) and other low frequency oscillations from sources other than the brain (Pinti *et al*., 2019). In general, fluctuations due to neural activity typically appear in the frequency range around 0.1 Hz. When analysing waveforms, a low pass filter is used to focus on signals below the heart rate variability band, and a high pass filter is used to remove low frequency components. Note that filtering is not required if only time-frequency analysis (e.g., wavelet transform coherence) is to be performed.

The artefact that cannot be eliminated by the above methods is skin blood flow. Since fNIRS measures brain activity by placing the optodes on the scalp, the skin blood flow always contaminates the signals. There are several statistical removal methods, which exploits the distinct statistical properties of skin blood flow compared to cortical activity (Zhang *et al*, 2005; Kohno *et al*., 2007; Yamada *et al*, 2012; Sato *et al*, 2016).

When assessing synchrony using correlation and wavelet transform coherence, prewhitening is recommended to remove serial correlation caused by haemodynamics in fNIRS time series. This process reduces false positives and facilitates more accurate identification of within-brain and between-brain synchrony (Santosa *et al*., 2017). When considering inter-brain synchrony between participants with different haemodynamics (e.g., adults and infants), pre-whitening can eliminate bias caused by such differences (Morimoto and Minagawa, 2022).

#### Measure of inter-brain synchrony

Two of the most popular measures (i.e., wavelet transform coherence and Pearson’s correlation coefficient) and an information metric (i.e., phase mutual information) for quantifying synchrony are summarized below.

##### Wavelet transform coherence

Wavelet transform coherence (WTC) can analyse the relationship between two signals in both time and frequency domains. When a continuous wavelet transform (CWT) of signal *X* is *W_X_*, a wavelet power spectrum is defined as the squared magnitude of *W_X_* and calculated as follows,

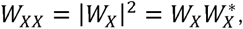

where * indicates the complex conjugate. A cross-wavelet transform of two signals, *X* and *Y*, is

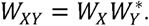

WTC is defined as the squared modulus of the smoothed cross-wavelet transform normalised by the product of the smoothed wavelet power spectra of each time series,

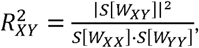

where *S* denotes a smoothing operator in both time and frequency. As the fNIRS signal is non-stationary, wavelet-based quantification is preferred to Fourier-based quantification (e.g., magnitude-squared coherence). Cui *et al*. first introduced WTC to assess neural synchrony between two participants during a cooperative tapping task (Cui *et al*., 2012). They calculated the average WTC within the task-relevant frequency bands during each task condition and converted it to Fisher’s *z*-statistic to perform a one-sample *t*-test. This procedure is a standard analytical flow of WTC. However, it should be noted that *z*-transform can unnecessarily distort the distribution of the average WTCs due to its non-negativity, as the lower and upper limits are 0 and 1, respectively.

##### Correlation coefficient

Pearson’s correlation coefficient (CC) is the simplest quantification of the linear relationship between two signals. The definition of CC is the covariance of the two variables divided by the product of their standard deviations,

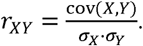

CC has been used as a standard measure of functional connectivity in fMRI studies (Misaki *et al*., 2021). The use of CC for hyperscanning analysis assumes that signals from interacting brains show simultaneous variations. This assumption is stricter than that of WTC. Although there are many variations of CC that extend its statistical properties, the original CC is still used in many hyperscanning studies (see Hakim *et al*., 2023). By introducing a sliding-window approach, CC can be extended to measure temporal-dynamic relationships between two signals. This extension, the sliding window correlation coefficient (swCC), has been the most popular approach to evaluate dynamic functional connectivity for fMRI signals (Hutchison *et al*., 2013). Note that there are multiple parameters for the sliding window (e.g., window function, duration, step size) and they directly affect the outcomes (Shakil *et al*., 2016; Mokhtari *et al*., 2019).

##### Mutual information

Mutual information (MI) is a non-directional, model-free measure of the amount of information between two random variables. It quantifies the reduction in uncertainty about one variable given knowledge of the other. MI has not been used for fNIRS signal analysis until recently (Sun *et al*., 2023).

The definition of MI between two signals is the Kullback-Leibler divergence between the joint probability distribution and the product of the marginal distributions as follows,

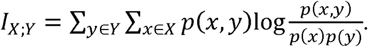

MI is always non-negative and symmetric with respect to the variables *X* and *Y*. When signals follow Gaussian distributions, relationship between MI and CC is as follows,

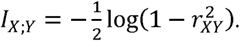

However, unlike CC, MI can assess non-linear relationships (Hlinka *et al*., 2011). MI can also be easily extended to time-series analysis by introducing the sliding-window approach (sliding window mutual information).

#### Behavioural association

For interactive social neuroscience, revealing the relationship between synchrony and behaviour is the most fundamental part. For static measures, the most straightforward approach is to examine the conditional differences in synchrony using statistical tests (e.g., group condition vs individual condition; typical development vs autism spectrum disorder). Direct comparisons of time-averaged coherence should be treated with caution due to the potential bias introduced by sample size (Maris *et al*., 2007; Bastos and Schoffelen, 2016). To assess the relationship with a scaled variable (e.g., questionnaire score), Peason’s correlation coefficient or Spearman’s rank correlation coefficient is often used. Linear regression analysis is also available if synchrony can be modelled by the linear combination of variables. When the measure is not normally distributed, use of a generalised linear model (GLM) is recommended.

For dynamic measures, modelling the time-varying relationship between the measure and behaviours is necessary. Despite its importance, few studies have examined such analyses. Liu *et al*. applied a linear model to WTC of task-relevant frequency bands (i.e., 0.08–0.04 Hz) (Liu *et al*., 2016). They constructed a boxcar shape regressor for each task condition and compared the estimated coefficients between conditions. Xu *et al*. evaluated social event-related changes in WTC using GLM analysis (Xu *et al*., 2023). They used an event-related box-car shape regressor with 5 seconds delay and assumed a gamma distribution as an error distribution. Since the participants freely interacted with each other, they did not focus on task-relevant frequency bands but the frequency bands of 0.1–0.03 Hz. Regressors in both studies, however, may not be optimal because they did not stand on the temporal properties of WTC; WTC change at the low-frequency band does not follow a boxcar-like shape. It should be emphasised that by adjusting the temporal and frequency characteristics of the WTC in the regressors, the detection of behaviourally relevant synchrony is expected to increase.

### Simulation settings

In this section, we performed simulations to examine the relationship between several static and dynamic measures for inter-brain synchrony listed in Table 1. We also conducted simulations of a WTC-GLM analysis to further explore the behavioural association with WTC. All signal processing and statistical analyses were performed using Matlab 2021a (The MathWorks, Natick, Massachusetts, USA).

#### Static measures

We conducted simulations using a resting-state data covering broad brain areas from a single participant because hyperscanning data are generally independent and not synchronised. We expected that such a dataset would exhibit different correlation patterns and be suitable for evaluating the differences between the measures. The data used were obtained from the MULDS dataset, collected by investigators at ATR, Japan, with funding support from the National Institute of Information and Communications Technology, Japan. The data include resting-state activities from 106 channels over the whole head, recorded over a period of 617.75 seconds at a sampling rate of 4 Hz.

For preprocessing, we applied haemodynamic modality separation (Yamada *et al*., 2012) to the converted Δoxy-Hb and Δdeoxy-Hb signals to remove systemic physiological noise, including motion artefacts and skin blood flow. A wavelet-MDL filter (Jang *et al*., 2009) was then applied to eliminate the low-frequency trend. The signals were prewhitened with an AR (40) filter (i.e., 10 s) to remove serial correlation. Only the Δoxy-Hb signals, excluding the first 10 seconds, were used for the following analyses.

WTCs were calculated with an analytical Morlet wavelet kernel and complex Gaussian wavelet kernels (first and second order derivatives). The number of voices was set to 12. In order to obtain a sufficient sample length for averaging, we focused on frequency bands above 0.03 Hz. All WTCs in each frequency band were averaged over time, except for the cone of influence (tWTC); we averaged the smoothed wavelet cross-spectrum along the time domain and then calculated its absolute value (Zhang *et al*., 2020). The average value of tWTC within 0.08–0.03 Hz was also calculated (tsWTC).

CC was calculated for the additional band-passed signals at 0.08–0.009 Hz through the third-order low-pass and the fifth-order high-pass Butterworth filter (Novi *et al*., 2016).

For mutual information, several implementations exist (e.g., Kraskov *et al*., 2004). In this simulation, we used phase mutual information (pMI; Palus, 1997) due to its computational simplicity. From the band-passed signals for CC, we obtained the instantaneous phase using the Hilbert transform. pMI was calculated using the whole binned phase sequences (Scott, 1992).

We calculated each measure for randomly chosen 1000 channel pairs.

#### Dynamic measures

We compared several dynamic measures for the resting-state dataset in the same way as the static measures. We selected WTC with an analytical Morlet wavelet kernel, swCC, and the sliding window extension of pMI (swpMI) for this simulation. For the sliding window measures, we used a simple rectangular window, two types of length settings (i.e., 30 seconds and 60 seconds), and the minimum step size (i.e., one sample).

#### Behavioural association with WTC

As described in the previous section, WTC-GLM analyses in the previous studies were not optimised for the characteristics of WTC. To obtain a regressor suitable to the non-linear response, we simulated the ideal WTC from the synchronised behavioural events of two participants.

Here, we assumed that inter-brain synchronised neural events are caused by sharing a social signal between two participants (i.e., the Social Signal Model in Morimoto and Minagawa, 2022). This simple assumption allows us to simulate ideal fNIRS response sequence pairs using boxcar-modelled neural events and the convolution of a haemodynamic response function (Zhang *et al*., 2020). By adding a different resting-state signal to each sequence as baseline noise, we obtained a synchronised fNIRS signal pair. We prepared a combination of a target WTC (i.e., the synthesised-target WTC) and 100 of *synthesised* WTCs (i.e., the synthesised-*synthesised* WTC) to construct a regressor. An ideal WTC was obtained by averaging the *synthesised* WTCs for each time step and frequency bands. The target WTC within the frequency bands of interest was modelled by a GLM with the regressors including the corresponding ideal WTC and a constant term. The error distribution and link function in the GLM were a gamma distribution and log-link function, respectively.

We used the resting-state data from an open dataset (von Lühmann *et al*., 2020) as baseline noise. This dataset includes 5 minutes of 26-channel resting-state data from the occipital area of 14 participants and 10 minutes of 48-channel resting-state data from the fronto-parietal area of another 14 participants. In addition, the dataset provides a synthetic haemodynamic response (HR) of Δoxy-Hb generated by a gamma function. The total length of the HR was 17.6 s, including a short blank at the end. Since the peak amplitude of the HR was adjusted to match the resting-state data, it is straightforward to estimate the performance of the proposed method in a realistic situation. Thus, we used the HR as an alternative to the convolutional model described above. We used the first 5 minutes of each participant’s data for the subsequent simulation. Since one participant had less than 5 minutes of data, only data from 27 participants were utilised. All data was resampled at 10 Hz from the original 50 Hz sampling rate.

We randomly generated 5-minute synthetic Δoxy-Hb sequences at a sampling rate of 10 Hz, with the probability of an HR event set to 0.0025. As we did not allow the HR to overlap, one HR was expected every 57.6 s. After adding the resting-state data, we performed preprocessing in the same manner as for the simulation of the static measures. WTCs were calculated using an analytical Morlet wavelet kernel. We estimated the coefficients β of the GLM and collected them repeatedly for 100 different synthetic Δoxy-Hb sequences. We further estimated βs with different HR amplitude ratios from the original (0.1, 0.2, ·, 0.9) and without HRs (i.e., resting-state data only). The β from the constant term was expected to reflect the bias of WTC caused by fluctuations from baseline activity. Thus, we only focused on the βs corresponding to the ideal WTCs. To present the performance of our method, we assessed the effect size between the βs with HRs and the βs without HRs.

We also generated additional 5-minute sequences to determine the frequency bands to be used in the GLM analyses. In these sequences, only a single HR event arises 150 seconds after the beginning of the sequence. We synthesised 300 pairs of the Δoxy-Hb sequences for calculating target WTCs (congruent HRs) and 100 pairs for constructing a regressor. For each frequency band, we calculated the Fisher’s *z*-transformed Pearson’s correlation coefficient between the regressor and the target WTCs. In addition, we calculated the z-values from the WTC with the HR onset ranging from 125 seconds to 175 seconds at 5-second intervals, excluding 150 seconds (incongruent HRs), and from the WTC without HRs (baseline).

Furthermore, to demonstrate the advantage of the WTC-GLM analysis over a single-signal GLM analysis (i.e., not the coherence of two signals), we performed a simulation of GLM with the wavelet power spectra (CWT-GLM) in the same manner as the WTC.

### Simulation Results

#### Static measures

The relationships between the typical static measures are presented in Figure 1 and Figure 2. CC was squared for direct comparison with other measures (sqCC). All combinations of the static measures showed a positive rank correlation. There was a higher correlation between tsWTC and pMI (*ρ* = 0.69) and between sqCC and pMI (*ρ* = 0.61) than between tsWTC and sqCC (*ρ* = 0.38). Correlations between tWTC and pMI were also higher than between tWTC and sqCC in the practical frequency bands (<0.2 Hz) (Figure 2). These results indicate that there was a large difference in measured synchrony between WTC and CC in our simulation settings. pMI had intermediate properties between WTC and CC.

**Figure 1.**
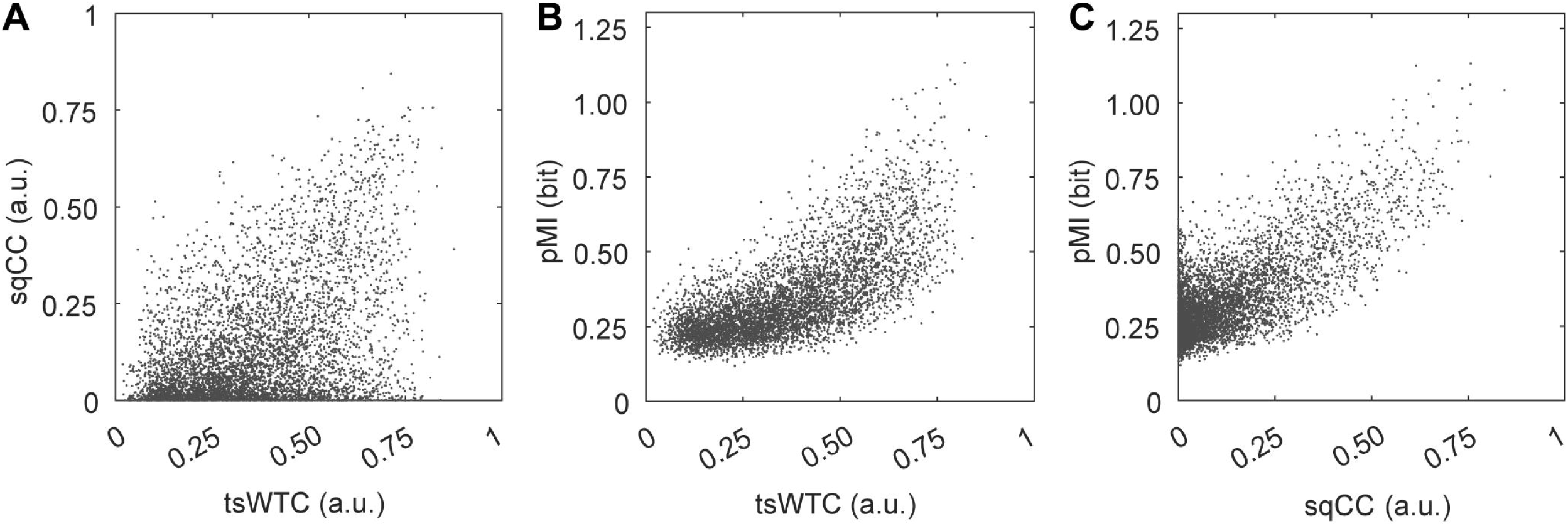
Relationship between static measures. Scatterplots between (A) tsWTC and sqCC, (B) tsWTC and pMI, and (C) sqCC and pMI. Each dot represents a pair of channels. See Figure S1 for comparisons with CC.

**Figure 2.**
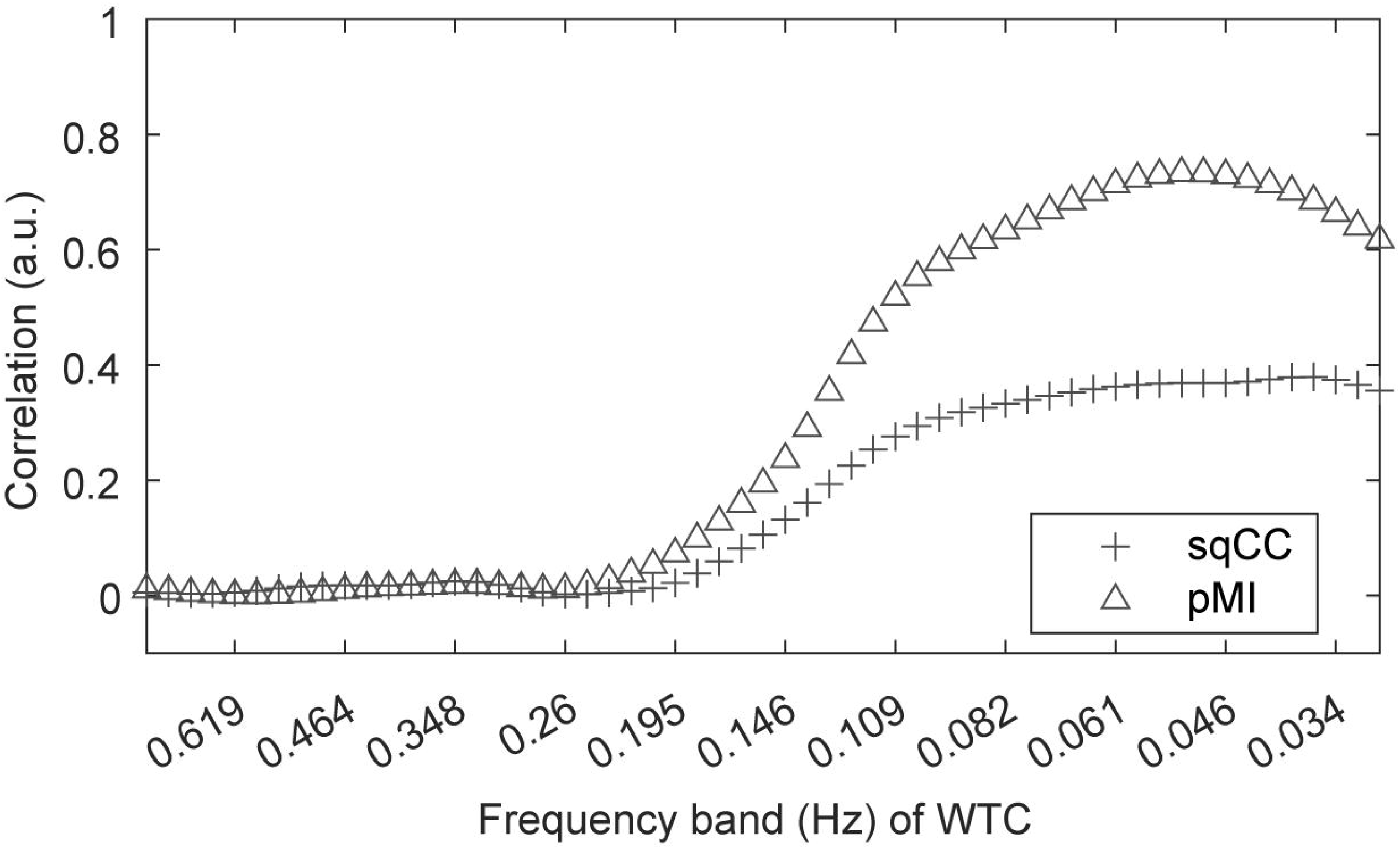
Correlation between tWTC and other static measures. Spearman’s rank correlation coefficients (p) are shown along the frequency bands of the tWTC. Plus signs and triangles indicate sqCC and pMI, respectively.

For further investigation, we compared the influence of prewhitening in the preprocessing on each static measure. Prewhitening had a greater effect on CC, while having less effect on tWTC (Figure S2). Interestingly, pMI showed a high correlation with tWTC regardless of prewhitening, whereas CC did not (Figure 2; Figure S3).

The influence of the wavelet kernel on tWTC is illustrated in Figure S4, with a greater impact observed in the lower frequency bands. The difference in the derivative order of the complex Gaussian kernel did not significantly affect tWTC.

#### Dynamic measures

The relationships between the WTC of each frequency band and other dynamic measures are shown in Figure 3. The correlation pattern illustrates the effect of the sliding window length. For both swCC and swpMI, the peak correlation with the WTC occurs around the frequency corresponding to the reciprocal of half the sliding window length. Compared to the case without prewhitening (Figure S5), prewhitening restores the correlation of the two measures of the 30-second sliding window.

**Figure 3.**
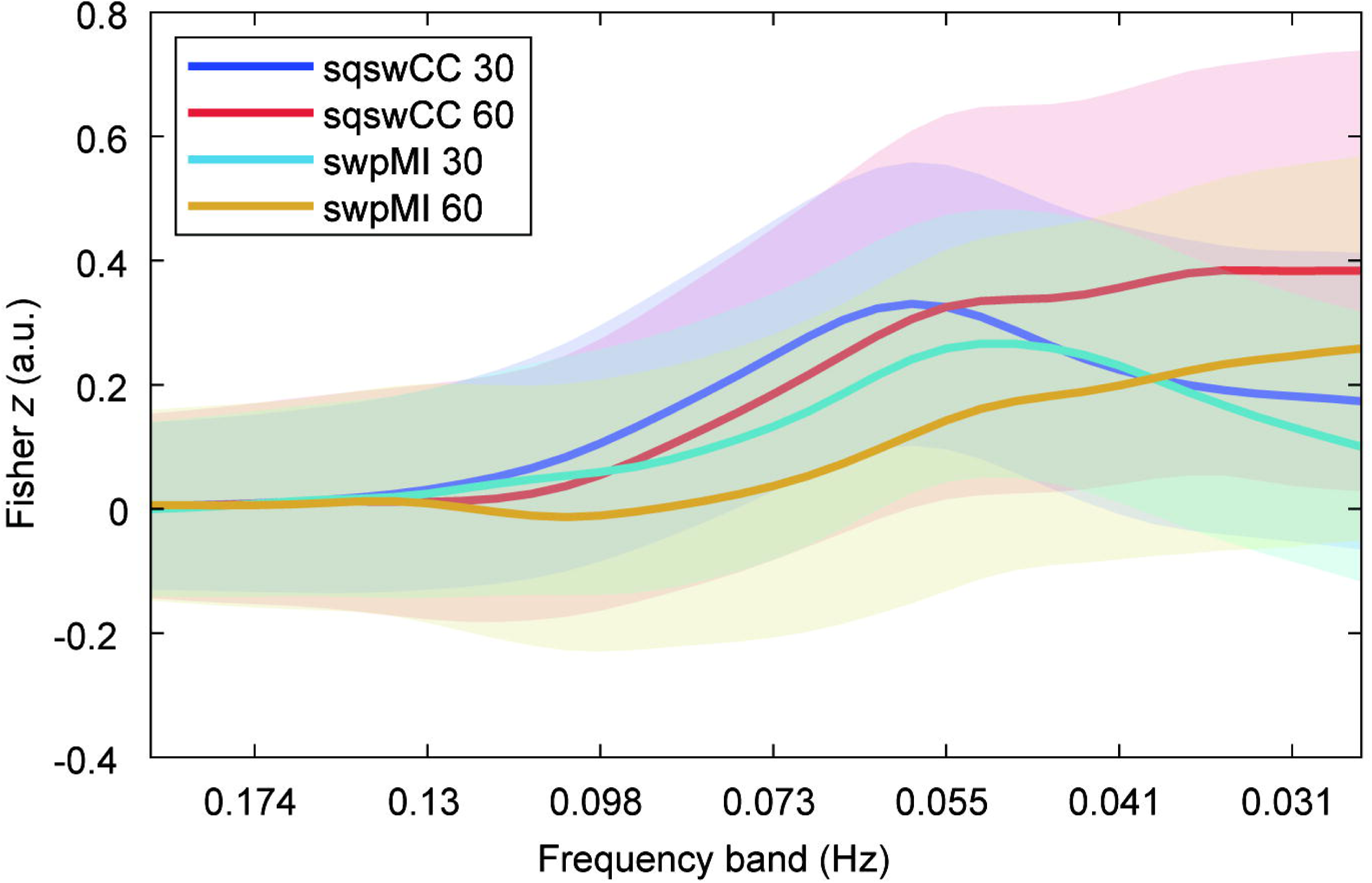
Correlation between WTC and other dynamic measures. The mean (bold line) and standard deviation (shaded area) of the *z*-transformed rank correlation coefficient were plotted along the frequency bands of WTC (sqswCC: squared sliding window CC, swpMI: sliding window pMI). The numbers in the legend indicate the length of the sliding window. For results without prewhitening, see Figure S5.

The relationships between swCC and swpMI are shown in Figure 4. Their correlation was higher for the 60-second sliding window than for the 30-second sliding window. Prewhitening had a small impact on this relationship (see Figure S6).

**Figure 4.**
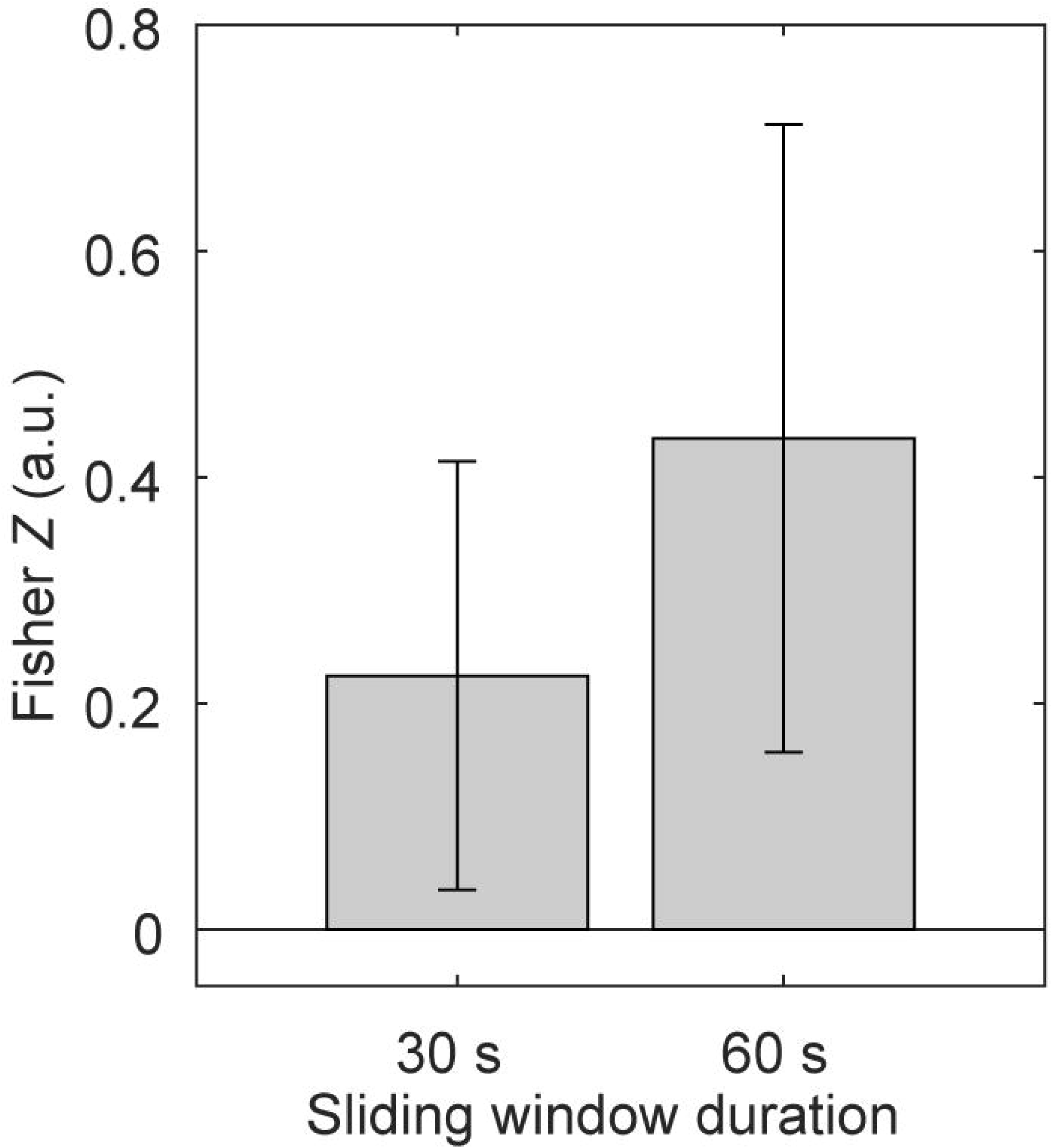
Correlation between swCC and swpMI. The mean and standard deviation of the z-transformed rank correlation were illustrated for each sliding window length. For results without prewhitening, see Figure S6.

#### Behavioural association with WTC

As has already been mentioned, determining the adequate regressor is crucial for the WTC-GLM. In this section, we propose a new method for it, which will be assessed and validated by simulations. First, the frequency bands for the GLM regression analyses were determined using the single HR simulation. Examples of the WTCs and corresponding regressors are shown in Figure 5. The amplitude of the WTC increased before the onset of the HR. In the lower frequency bands, it appeared difficult to distinguish small differences in HR onset due to the lower frequency shape of the regressor (Figure 5B). The correlation between the target WTC and the estimated regressor is illustrated in Figure 6. There are significant differences in the *z*-scored correlation between the congruent condition and the baseline condition, but smaller differences between the congruent condition and the incongruent condition. To accurately detect inter-brain synchrony caused by simultaneous HR pairs, it is probably essential to model not only the low-frequency bands (e.g., below 0.1 Hz) but also higher frequency bands. Based on these results, we utilized 0.2–0.07 Hz for the following WTC-GLM and CWT-GLM analyses. For comparison, we also calculated GLMs using data in the 0.1–0.07 Hz range.

**Figure 5.**
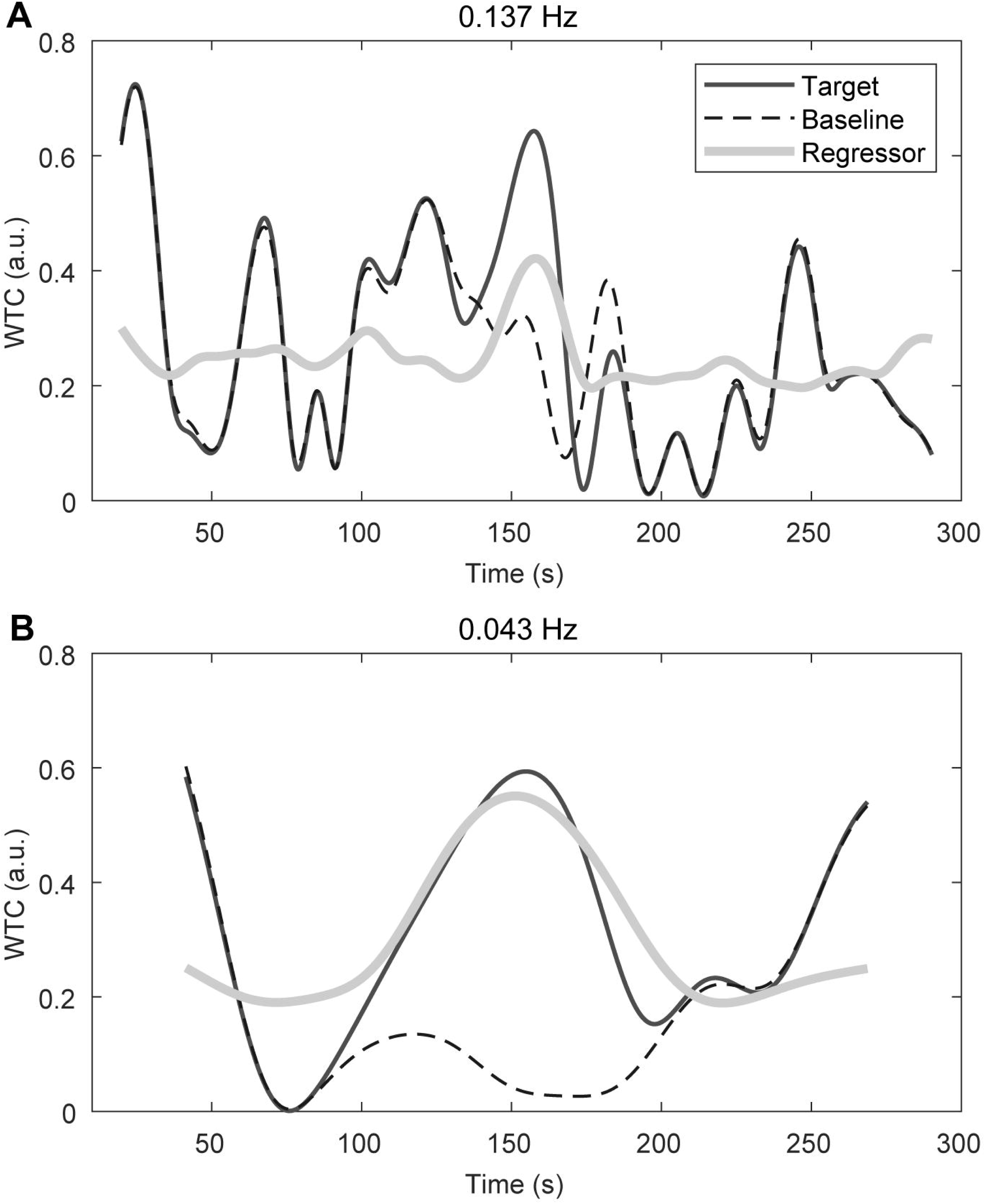
Examples of the regressors for WTC-GLM. The target WTC, the baseline activity (resting-state noise only), and the regressor are plotted as black solid, dashed, and gray solid lines, respectively, for the frequency bands of (A) 0.137 Hz and (B) 0.043 Hz. Note that the onset of the HR was set at 150 seconds from the beginning.

**Figure 6.**
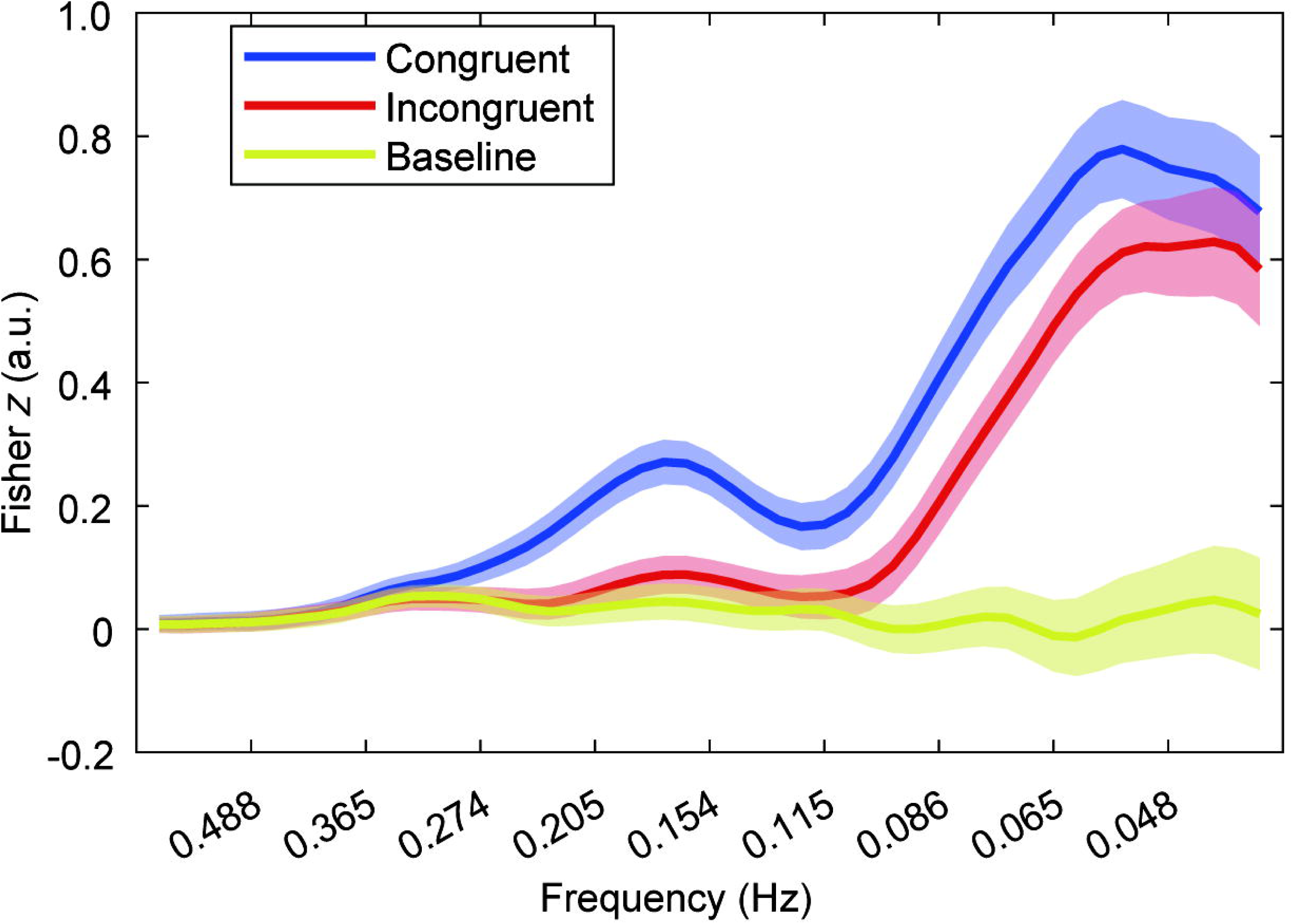
Correlation between the regressor and the target WTC. The *z*-scored rank correlations for three types of target WTC (congruent HRs, incongruent HRs, and baseline) are illustrated. The line and shaded area represent the mean and 95% confidence interval (CI) of each frequency band.

The distributions of coefficients, β, for WTC-GLM and CWT-GLM are presented in Figure S7 as an example. We calculated Cohen’s *d* to assess the difference between the distribution in the congruent condition and that in the baseline condition. WTC-GLM had higher *d*-values than CWT-GLM, and it increased as the amplitude of the HRs increased (Figure 7). The *d*-values were higher than those obtained from data in the 0.1–0.07 Hz range. Furthermore, the *d*-values obtained by comparing the congruent and incongruent conditions also showed similar patterns, except when the amplitude ratio was less than 0.3 (Figure S8). These results suggest that WTC-GLM can more precisely detect simultaneous HRs compared to CWT-GLM.

**Figure 7.**
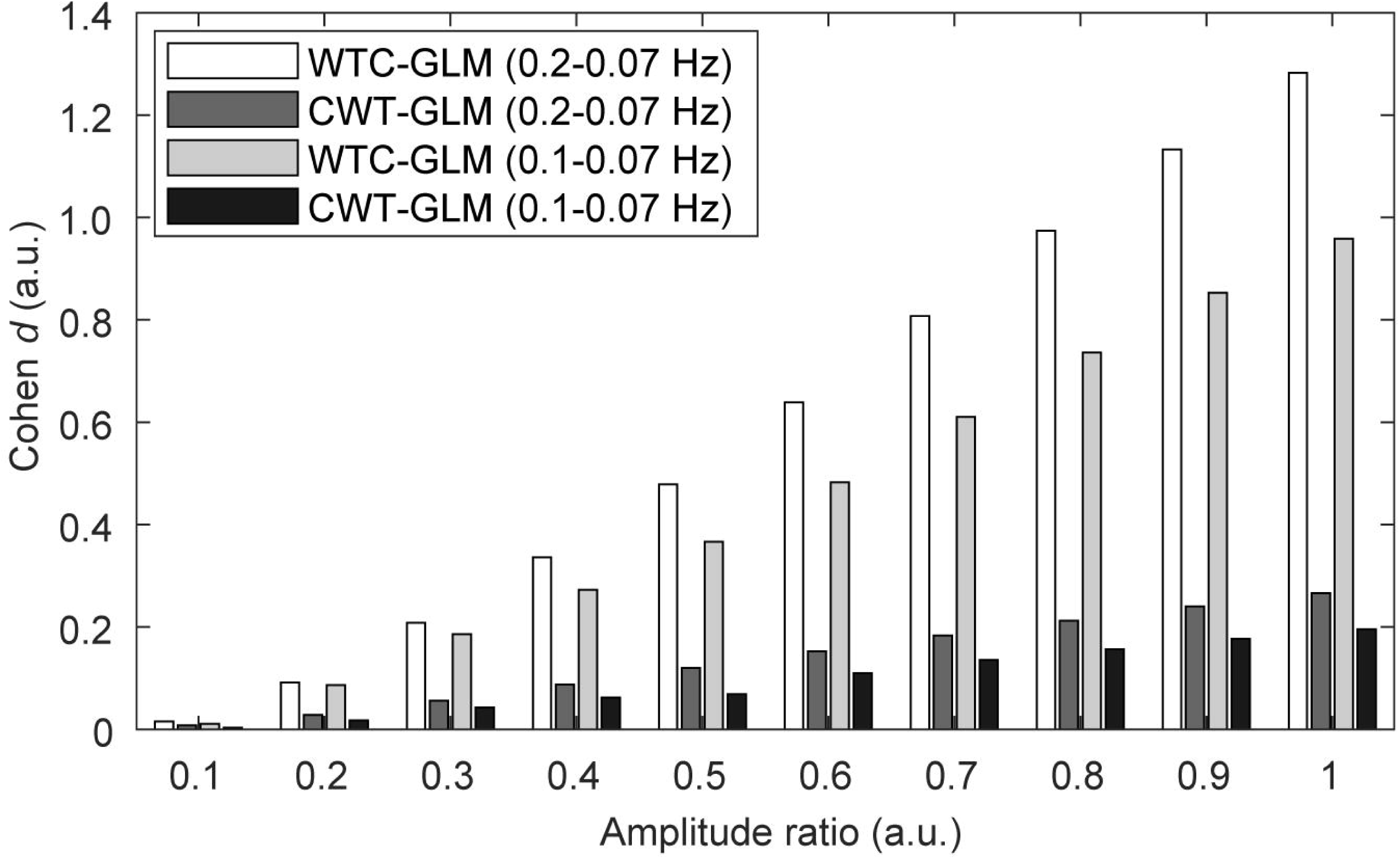
Detection performance of the haemodynamic response as a function of its amplitude. Each bar represents Cohen’s *d* between the beta values of the congruent condition and those of the baseline condition. Results are shown for the combinations of WTC-GLM and CWT-GLM within the frequency ranges of 0.2–0.07 Hz and 0.1–0.07 Hz.

## Discussion

In the present study, we performed simulations of several measures of inter-brain synchrony. Using the resting-state data, we examined the characteristic differences between static and dynamic measures. Simulation results indicated that WTC and CC highlighted different aspects of inter-brain synchrony in fNIRS signals. When averaging WTC over time series and frequency bands of interest (i.e., tsWTC), its properties are more similar to pMI than to CC. However, this tendency was not observed for WTC as a dynamic measure. These results suggest that, despite the simple averaging process, the aspect of quantifying synchrony changes. We have also proposed a new method to update the WTC-GLM analysis, which resulted in increasing its sensitivity to simultaneous haemodynamic responses between dyads. Simulation results showed that a more realistic shape of the regressor and modelling of a wider range of frequency bands (i.e., 0.2–0.07 Hz) was effective in detecting the associated inter-brain synchrony.

Although WTC is a standard measure for analysing fNIRS hyperscanning data, its properties for fNIRS signals have not been sufficiently explored (Zhang *et al*., 2020). While CC assumes a strict linear relationship between two signals, WTC can capture non-linear relationships. In addition, WTC’s flexibility allows it to assess the out-of-phase relationship between two signals (Hakim *et al*., 2023). These differences are likely to result in a significant difference in the measured synchrony between WTC and CC (see Figure 1A and Figure 2).

Instantaneousness is another key feature of WTC that explains its relationship with pMI. pMI is also a non-linear measure of synchrony, but its correlation with WTC changes between use as a static measure and use as a dynamic measure (see Figures 2 and 3). This difference can be explained by their definitions: while WTC evaluates the local (i.e., instantaneous) relationship, pMI evaluates the temporally averaged relationship over a longer period to calculate the probability. Therefore, the correlations between pMI and tWTC, as well as between pMI and tsWTC, were higher than expected based on the correlation between swpMI and WTC. The instantaneous nature of WTC also affects its difference from CC. It should be noted that this feature offers potential to utilize for behavioural association analysis.

Although predicting the dynamic shape of synchrony in WTC is difficult due to the non-linearity of WTC and fNIRS haemodynamic signals, we proposed a solution. To achieve a more accurate WTC-GLM analysis, we adopted a model-based approach; the shape of the regressor was optimised by repeatedly generating synthesised signal pairs with different types of resting noise and calculating the average WTC. Our method correctly detected the amplitude change of HRs (Figure 7) and achieved significantly high performance in discriminating incongruent (i.e., unpredicted) inter-brain synchrony (Cohen’s d > 1, when the amplitude of HRs was normal). The WTC-GLM without higher frequency bands (> 0.1 Hz) (used in Liu *et al*., 2016 and Xu *et al*., 2023) showed lower *d*-values. The inclusion of these higher frequency bands seems to be important for achieving accurate performance. However, our simulation settings of WTC-GLM did not cover various types of task designs but focused solely on a 5-minute free interaction task, with the occurrence of a social event approximately once every minute on average. For example, Pinti *et al*. reported a 2-second delay in haemodynamic responses between the informer and the guesser as a result of GLM analysis (Pinti *et al*., 2021). In our settings, we assumed simultaneous HRs (i.e., no delay). The optimal settings and performance limits of WTC-GLM would depend on the participants’ behaviour and the task design. We strongly recommend performing simulations prior to conducting fNIRS experiments, taking advantage of a model-based approach.

The difference in performance between WTC-GLM and CWT-GLM demonstrated the effectiveness of WTC’s robustness to noise. We suspect that the calculation of the WTC provides a denoising effect similar to that of trial averaging. Since trial averaging is not possible for the free interaction task, the denoising effect of WTC has the potential to more accurately assess task-related changes in fNIRS signals, not only for the assessment of inter-brain synchrony.

Although the present study has only focused on non-directional inter-brain synchrony, causality is also important for assessing the interaction between two brains (see Table 1). Granger causality is a popular measure of causality and has been used in an fNIRS hyperscanning study (Ono *et al*., 2022). Transfer entropy has also the potential to assess complex interactions in hyperscanning data, since it does not assume linearity as opposed to Granger causality (Wang and Chen, 2020; Wang, Chen *et al*., 2022). Additionally, several approaches to causality have utilised extended synchrony measures, such as cross-correlation (Bizzego *et al*., 2022) and cross-generalised linear models (Pinti *et al*., 2021). Despite its importance, there have been limited studies attempting to approach inter-brain causality. This is because static measures of inter-brain synchrony can only be used in limited situations (e.g., fixed leader-follower relationships). To analyse the dynamic inter-brain relationship during completely free interaction, future studies should develop more robust dynamic measures of causality (e.g., sliding window approach).

## Conclusion

Focusing on three different measures of inter-brain synchrony, our simulations revealed the relationships between them. In general, WTC was found to quantify synchrony with a broader definition due to its advantages of non-linearity, instantaneousness, and robustness. Conversely, pMI exhibited similar performance to the temporally averaged WTC, while CC quantified synchrony with a more limited definition. Additionally, we proposed a new method of GLM regression analysis for WTC to investigate the behavioural association of inter-brain synchrony. Our simulations validated the WTC-GLM analysis, enabling us to more precisely quantify the relationship between brain synchrony and behaviour than previous methods. Our WTC-GLM results also suggested the potential of WTC to be a robust metric for analysing free interaction data.

Recently, interest in the neural mechanisms supporting social interaction has been rapidly expanding, and fNIRS, with better spatial resolution than EEG, has significant potential to reveal new evidence by determining localised brain functions in natural interactions. Since real-life social interaction is dynamic rather than static, behavioural modelling and its associations with inter-brain synchrony are becoming increasingly important. This simulation-based study, including the proposed GLM analysis, represents a step forward in enhancing research on social interaction.

## Data Availability Statements

The data underlying this article are available in Zenodo, at https://dx.doi.org/10.5281/zenodo.12741292. The datasets were derived from sources in the public domain: Multi-modal and multi-task human brain imaging dataset, https://bicr-resource.atr.jp/mulds/; Open Access Multimodal fNIRS Resting State Dataset With and Without Synthetic Hemodynamic Responses, https://www.nitrc.org/frs/?group_id=1071.

## <SUPPLEMENTAL Figures>

**Figure S1:** Relationship between static measures. Scatterplots between (A) CC and tsWTC, and (B) CC and phase MI. Each dot represents a pair of channels.

**Figure S2:** Influence of prewhitening on the static measures: (A) tsWTC (average tWTC over 0.08–0.03 Hz), (B) CC, (C) squared CC, (D) pMI, (E) tWTC at 0.310 Hz, (F) tWTC at 0.077 Hz, (G) tWTC at 0.49 Hz, and (H) tWTC at 0.031 Hz, respectively. Each dot represents a pair of channels. The dashed line indicates the line of equality.

**Figure S3:** Correlation between tWTC and other static measures, without prewhitening. Spearman’s rank correlation coefficients (p) are shown along the frequency bands of the tAWTC. Pluss sign and triangles indicate sqCC and pMI, respectively.

**Figure S4:** Influence of the difference of wavelet kernels on tWTC. The left, middle, and right columns show the relationship between analytical Morlet and complex Gaussian (first order), between analytical Morlet and complex Gaussian (second order), and between complex Gaussian of first and second order, respectively. The dashed line indicates the line of equality. (amor: analytical Morlet; cgau1: complex Gaussian of first order derivative; cgau2: complex Gaussian of second order derivative)

**Figure S5:** Correlation between WTC and other dynamic measures, without prewhitening. The mean (bold line) and standard deviation (shaded area) of the *z*-transformed rank correlation coefficient were plotted along the frequency bands of WTC (sqswCC: squared sliding window CC, swpMI: sliding window pMI). The numbers in the legend indicate the length of the sliding window.

**Figure S6:** Correlation between swCC and swpMI, without prewhitening. The mean and standard deviation of the *z*-transformed rank correlation were illustrated for each sliding window length.

**Figure S7:** Example of the distribution of beta coefficients. Histograms of beta values for (A) WTC-GLM and (B) CWT-GLM when the target frequency range was 0.2–0.07 Hz and the amplitude ratio was 1 are shown. Blue and red indicate the beta values from the congruent and baseline conditions, respectively.

**Figure S8:** Discrimination performance of the haemodynamic response as a function of its amplitude. Each bar represents Cohen’s *d* between the beta values of the congruent condition and those of the incongruent condition. Results are shown for the combinations of WTC-GLM and CWT-GLM within the frequency ranges of 0.2–0.07 Hz and 0.1–0.07 Hz.

